# Chromosome level draft genomes of the fall armyworm, *Spodoptera frugiperda* (Lepidoptera: Noctuidae), an alien invasive pest in China

**DOI:** 10.1101/671560

**Authors:** Huan Liu, Tianming Lan, Dongming Fang, Furong Gui, Hongli Wang, Wei Guo, Xiaofang Cheng, Yue Chang, Shuqi He, Lihua Lyu, Sunil Kumar Sahu, Le Cheng, Haimeng Li, Ping Liu, Guangyi Fan, Tongxian Liu, Ruoshi Hao, Haorong Lu, Bin Chen, Shusheng Zhu, Zhihui Lu, Fangneng Huang, Wei Dong, Yang Dong, Le Kang, Huanming Yang, Jun Sheng, Youyong Zhu, Xin Liu

**Author notes:** These authors contributed equally to this work (Huan Liu, Tianming Lan, Dongming Fang, Furong Gui). Correspondence: Xin Liu, Youyong Zhu, or Jun Sheng.

## Abstract

The fall armyworm (FAW), *Spodoptera frugiperda* (J.E. Smith) is a severely destructive pest native to the Americas, but has now become an alien invasive pest in China, and causes significant economic loss. Therefore, in order to make effective management strategies, it is highly essential to understand genomic architecture and its genetic background. In this study, we assembled two chromosome scale genomes of the fall armyworm, representing one male and one female individual procured from Yunnan province of China. The genome sizes were identified as 542.42 Mb with N50 of 14.16 Mb, and 530.77 Mb with N50 of 14.89 Mb for the male and female FAW, respectively. We predicted about 22,201 genes in the male genome. We found the expansion of cytochrome P450 and glutathione s-transferase gene families, which are functionally related to the intensified detoxification and pesticides tolerance. Further population analyses of corn strain (C strain) and rice strain (R strain) revealed that the Chinese fall armyworm was most likely invaded from Africa. These strain information, genome features and possible invasion source described in this study will be extremely important for making effective strategies to manage the fall armyworms.

## 1 Introduction

It has been more than 100 years since the Fall armyworm (FAW), *Spodoptera frugiperda* (J.E. Smith) was reported to damage maize and other crops in the USA^1^. It is a severely destructive agricultural pest native to Americas which survives the whole year in the tropical and subtropical area from far south Argentina, Chile and La Pampa to far north Florida, Texas, Mexico and the Caribbean^2–5^. It cannot survive severe winters because of the lack of diapause. However, FAW has a remarkable capacity of long-distance migration, with which the FAW can fly over 100 km per night^6^. Each spring, it can migrate over 2000 km from the overwintering areas to reinvade more northern regions, even up to Canada^4,7,8^. Recently, FAW spread out from its native region and invaded into Africa in 2016 with the report in São Tomé, Bénin, Togo and Nigeria^9^. This invasion rapidly became widespread in the whole sub-Saharan Africa till October 2017^10,11^. Following this trend, FAW soon invaded into many Asian countries, including India, Yemen, Thailand, Myanmar and Sri Lanka in 2018^12–14^.

Early R *et al* (2018)^5^ forecasted that China is one of the most vulnerable countries of being invaded by the FAW according to the information on frequent commercial trade and passenger transportation between Africa and China. Half a year later after this forecast, in January 2019, International Plant Protection Convention (IPPC) Contact Point for China spotted FAW for the first time in Puer and Dehong city, Yunnan Province, China (https://www.ippc.int/en/news/first-detection-of-fall-armyworm-in-china/). This invasive pest has rapidly invaded many provinces in China by June 2019. Now, the FAW has been detected in large parts of the world (Figure 1).

**Figure 1.**
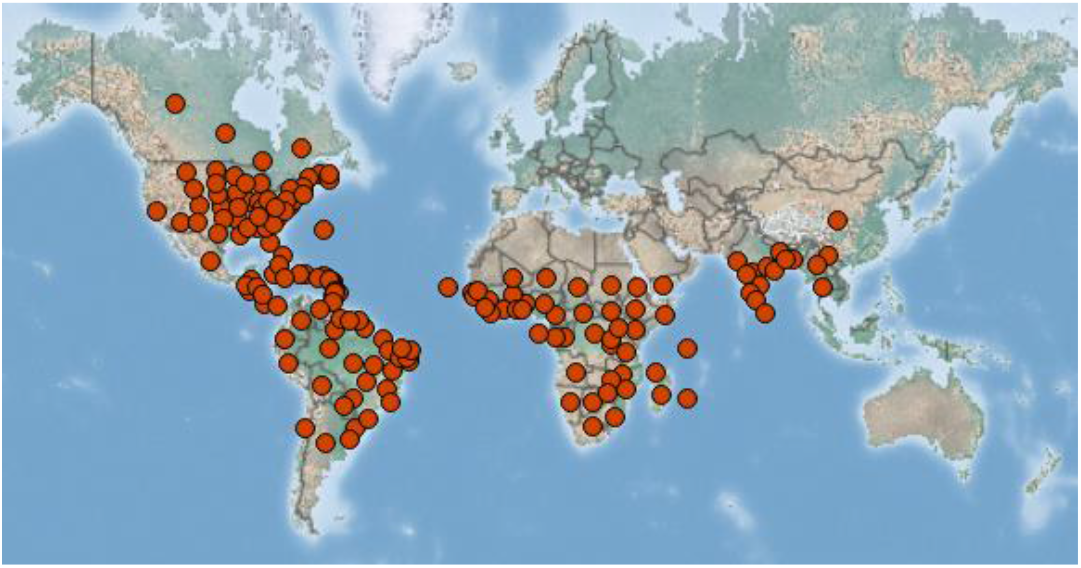
The distribution of the FAW all over the world^15^.

There are several hosts of FAW, which mainly includes 186 plant species from 42 families^16^. Although FAW is a highly polyphagous pest, graminaceous plants are their preferred hosts, such as maize, rice, sorghum and other major agricultural species. Especially, maize is most likely to be attacked than other plant species^9^. Maize is very important food security of many countries in Asia, Africa, and Latin America (https://maize.org/projects-cimmyt-and-iita-2/), and now are facing severe threats from the infestation of the FAW. The production of maize infested by the FAW can suffer yield loss of 40% to 72%, and in some plots 100% total has been reported^17–19^. The yield loss of maize can reach 8.3 to 20.6m tons per year in just 12 African countries without any control methods for FAW, according to the study conducted by Centre for Agriculture and Biosciences International (CABI)^20^. For Brazil alone, cost to control the FAW on maize is more than 600 million dollars per year^21^. The economic and the yield losses by FAW is a major concern worldwide.

The FAW is not a new species to science; it has been an herbivorous pest for many years. However, the mostly common used approach to mitigate the damage to crops is still the broad-spectrum insecticides^22^. The use of insecticides highly relies on the knowledge of farmers, but many farmers even do not know the name of the pest^23^. Lackof scientific guidance leads to the inappropriate use of pesticides. Besides, the effective management of FAW may require five sprays of pesticides per maize cycle^21^, and many smallholder farmers cannot afford the expensive costs for it, resulting in the use of low-quality products or low dose sprays^24^. Moreover, the FAW always hides inside the stem of maize, this makes the insecticide much less effective. Although many pesticides are less harmful to the environment and humans, all these factors can lead the sublethal effects, which possibly help the FAW to evolve resistance against the pesticides^25–27^. An effective compensating management for insecticides is the use of *Bacillus thuringiensis* (Bt) toxins produced by the by the bacterium.Bt plants have been proved fatal to many insect pests, including the FAW^28–30^. The Bt toxin provides much longer protection than insecticides and less harmful to the environment and humans. Although some research reported the resistance of FAW to Bt maize^29,30^, multiple genes or new gene with more or new Bt toxin expressed are still thought to have a good performance for resisting the FAW^29,30^. Biological control, including the, introduction of natural enemies and using companion cropping system^19,21,25,31,32^, is also an effective way to resist the FAW.

The FAW at least consist of two morphological identical but genetically distinct subpopulations, the corn strain (C strain) and rice strain (R strain)^33–35^. The two strains have their own preferable host plants. The C strain is preferentially associated with maize and other large grasses, but the C strain prefers rice and large grasses^36,37^. The C strain is subdivided into two subgroups, the FL-type and the TX-type^38,39^. The TX-type is distributed in most of the Americas, but the FL-type is only limited to Florida and the Caribbean^3,35,40,41^. Each strain has its strain specific physiological traits, leading to some strain-specific response to biological and chemical agents^36,37,42^. The C strain larvae are more tolerant than the R strain to the methyl parathion, cypermethrin, cypermethrin and ∂-endotoxin from transgenic Bt plant^43^. Therefore, the origin of FAW needs to be considered to make effective strategies to manage the FAW. The *Cytochrome c oxidase subunit I* (*COI*) and the *Triosephosphate isomerase* (*Tpi*) gene are also selected for identifying the subtype of the FAW, but these markers cannot always give the right identifications^44^.

The management of the FAW needs more detailed genetic information to help people know more about the FAW, to find more new genes to develop more effective Bt plants, and to more accurately identify the different strains for precision spraying of pesticides. Although several genomes of the FAW has been sequenced and assembled^45–47^, the assemblies were fragmented. Moreover, two genomes were assembled using the Sf21^45^ and Sf9^47^ cell line, respectively, the resource was unique that cannot well provide a comprehensive reference. In this study, we assembled two FAW genomes to the chromosome level using two samples (one male, one female) collected from Yunnan Province, China. We analyzed the subtype of the FAW invaded Yunnan province, and also discussed the possible resource of the invaded FAW in China. We are also screening expanded gene families to seek some key genes with the function of polyphagia and tolerance to insecticides. China is the second largest corn producer after the USA; therefore, it is urgent to select a series of methods to control the FAW. This study provided key information to help make strategies to manage the FAW in China.

## 2 Materials and Methods

### 2.1 Samples and treatments

We collected seven FAW samples, including four adults, two fifth-instar larvae and two sixth-instar larvae (Table 1). The four adult FAWs were collected from Yunnan Province, China, and the larvae were collected from Guangdong Province, China. One male and one female adult individual were used for genome sequencing and assembly (Figure 2). Two other adult FAWs were used to capture the conformation of chromosomes to perform the chromosome level genome assembly. One fifth-instar larva and one sixth-instar larva were subjected for the transcriptomic studies. One sixth-instar larva was used for whole genome sequencing with 5K and 300bp insert size. All samples were intestinal and ovarian, and thoroughly cleaned before performing DNA or RNA isolation.

**Table 1.**
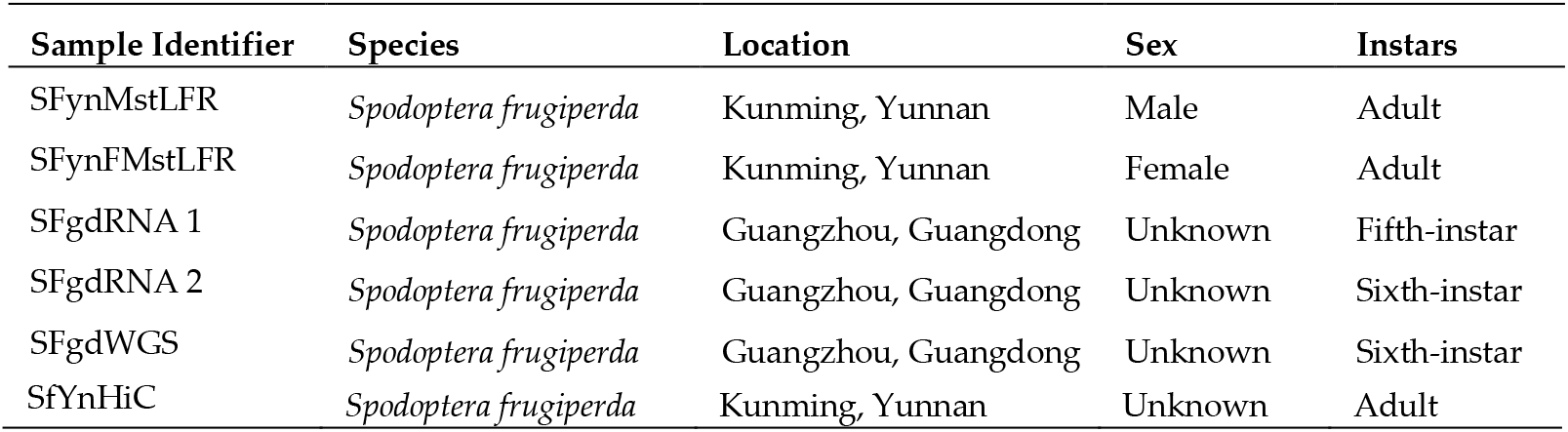
Samples used in this study

**Figure 2.**
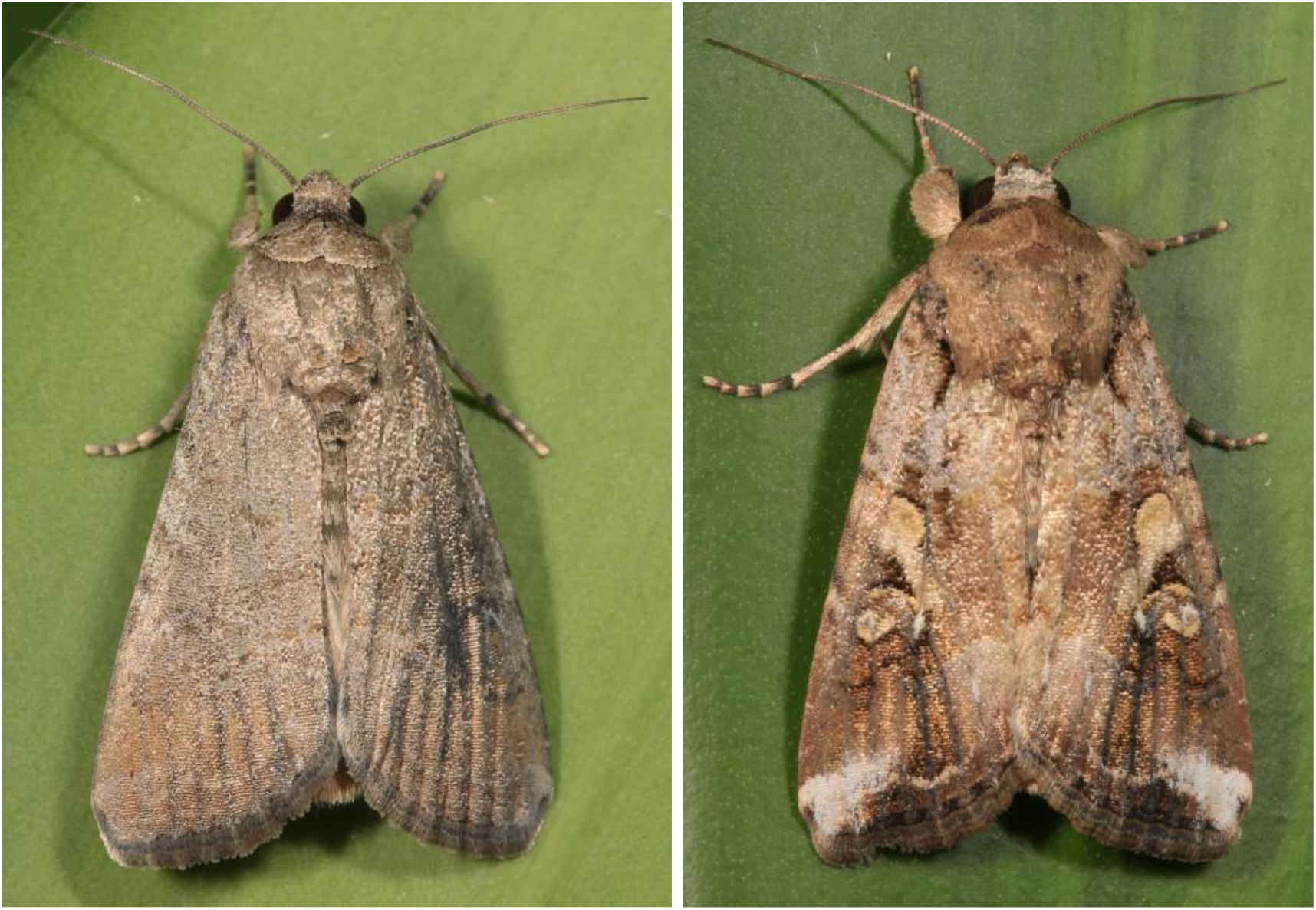
The two adult fall armyworm (FAWs) used for genome assembly. The left FAW is female, the right FAW is the male individual.

### 2.2 DNA isolation, library preparation, sequencing and genome assembly

The high molecular weight DNA was extracted using the separated muscle tissue by following the protocol recommended by Cheng et al (2018)^48^. We then used the single tube long fragment read (stLFR) technology^49^ to preparing the co-barcoding DNA libraries with the MGIEasy stLFR Library Prep Kit (lot number: 1000005622), and the libraries were then loaded on the sequencer for sequencing according to the protocol of MGISEQ-2000^50^. To testify the accuracy of genome assembly, we constructed a DNA library with a 5kb insert size and sequenced by BGISEQ-500 sequencer. To finally ligate the scaffolds to chromosomes, Hi-C technology^51^ was used to capture the conformation of chromosomes using another two adult individuals. The primary genome was assembled using the supernova (v2.1.1)^52^ software with the parameters --*maxreads*=*700M*. We filled gaps by use of GapCloser^53^ and GapCloser stLFR (unpublished method) with the default parametes. Finally, we performed the chromosome concatenation using the Hi-C generated data by 3d-DNA pipeline^54^.

### 2.3 Comparative genomics analysis

Identification of orthology and paralogy groups of *Spodoptera frugiperda* genes and other considered genomes were done using OrthoMCL^55^ methods on the all-versus-all BLASTP alignment (e-value, <1e-5). We constructed gene families for nine species including *Bombyx mori, Danaus plexippus, Drosophila melanogaster, Heliconius melpomene, Helicoverpa armigera, Manduca sexta, Plutella xylostella, Spodoptera litura and Spodoptera frugiperda*. The phylogenetic tree, including these nine species, was constructed using the combined set of all the single copy genes.

To search for homology, we compared protein-coding genes of *Spodoptera frugiperda* to that of other species using BLASTP with an E-value threshold of 1e-5. Base on the Whole-genome BLASTP and the genome annotation results, we detected the syntenic blocks using MCscan^56^. A region with at least five syntenic genes and no more than 15 gapped genes was defined as a syntenic block.

### 2.4 RNA isolation, transcriptome libraries preparation and sequencing

The RNA extraction kit (RNeasy Mini Kit, Qiagen) was used for the total RNA isolation. We performed the RNase-free agarose gel to check the contamination, and then the RNA integrity and purity were measured by Agilent 2100 Bioanalyzer system (Agilent, United States) and NanoDrop Spectrophotometer (THERMO, United States), respectively. The extracted RNA was fragmented into 200-400 bp and reverse transcribed to cDNA for library preparation. The libraries were prepared to follow the manufacturer’s instructions for the BGISEQ-500 sequencing platform. Pair-end 100 sequencing was performed on the BGISEQ-500 sequencer using the processed libraries.

### 2.4 Bioinformatics analysis for transcriptome data

Raw data were firstly processed using the Trimmomatic to filter the reads with adaptors and reads with the proportion of Ns and low-quality bases larger than 10% by SOAPfilter. Bridger software (v20141201) was used to *de novo* assemble the transcriptome, the reluctances were then removed by TGICL. The contigs were concatenated into scaffolds and further assembled to unigenes by clustering and removing redundancy.

FPKM was calculated to estimate the expression level of unigenes. In the study, the reads were mapped against the unigene library using Bowtie, and then unique mapped reads were selected for estimating the expression level by combining eXpress. Finally, DEG unigenes were selected with differential expression level with the parameter of FDR ≤ 0.01 and Fold Change ≥4.

### 2.5 Identifying the strains and the possible source of FAW invasion in China

We used the *Tpi* gene as a DNA maker to identify the strain of the FAW invaded into China. We identified the *Tpi* gene fragments from four FAWs, including two from Yunnan province (sequences were retrieved from the whole genome sequencing data) and two from Guangdong province (sequences were retrieved from the RNA-seq data). Eight sites (TpiE4-129, TpiE4-144, TpiE4-165, TpiE4-168, TpiE4-180, TpiE4-183, TpiE4-192, TpiE4-198) in the fourth exon of the *Tpi* gene were used for determining the strains and a possible source of the invaded FAW^57^. The fourth intron of the *Tpi* gene^57^ was used for constructing the phylogenetic tree using the PhyML (v3.0)^58^ software with the Maximum-Likelihood methods to assist in identifying the strains. Only two Tpi fourth introns were used for phylogenetic analysis because we cannot retrieve the introns from RNAseq data. 112 sequences of the fourth intron of the *Tpi* gene with strain information were downloaded from NCBI (Table S1). The sequence alignments were performed using the Clustal W^59^.

## 3 Results and discussion

### 3.1 Chromosome level genome assembly for two FAWs

Only one individual was used for genome sequencing for both the male and female genome assembly, which was less than the number used for genome assembly by Gouin A *et al* (2017)^46^. This maximally decreased the heterozygosity level. A total of 2μg and 1.6μg total genomic DNA with the average band size larger than 20 kb were isolated for stLFR libraries preparation from the male and female FAW, respectively. Two stLFR libraries with more than 30 million barcodes were constructed for running on the BGISEQ-500 sequencer. 110.93 Gb and 91.41 Gb high-quality reads were generated for the male and female individual, respectively (Table S2). The primary assembly sizes were 542.42 Mb and 530.77 Mb for the male and female individual. The scaffold N50 and N90 for the male individual were 507.12 Kb and 6.43 Kb, and that for the female were 528.27 Kb and 5.11 Kb, respectively (Table 2). After getting the confirmation from Hi-C sequencing, we finally concatenated the scaffolds to 31 chromosomes with the scaffold N50 of 14.16 Mb and 14.89 Mb for the male and female individual, respectively. Genomes sizes assembled in this study are in the range of the lepidoptera genome sizes at 246M to 809M^60^, but are larger than previously assembled FAW genomes^45–47^, probably due to the use of the new stLFR technology with the super high coverage and reads quality. This is the first time to assemble the genome of the FAW to the chromosome level, which will be in no doubt to accelerate the biological studies and making the effective strategy of pest managements.

**Table2.** Summary of the two Spodoptera frugiperda genome assemblies

| Methods | Statistics | SFynMstLFR |  |  |  | SFynFMstLFR |  |  |  |
| --- | --- | --- | --- | --- | --- | --- | --- | --- | --- |
|  |  | Scaffold<br>Length (bp) | Number | Contig<br>Length (bp) | Number | Scaffold<br>Length (bp) | Number | Contig<br>Length (bp) | Number |
| stLFR | N50 | 507,121 | 226 | 91,970 | 1,220 | 528,269 | 231 | 83,007 | 1,299 |
|  | Total length (bp) | 542,424,128 |  | 505,703,627 |  | 530,766,122 |  | 496,217,290 |  |
|  | Ratio of Ns (%) | 6.77 |  |  |  | 6.51 |  |  |  |
|  | GC content (%) | 36.52 |  |  |  | 36.57 |  |  |  |
| HiC | N50 | 14,162,803 | 16 | 91,970 | 1,220 | 14,883,732 | 13 | 124,992 | 813 |
|  | Total length (bp) | 543,659,128 |  | 505,703,627 |  | 531,931,622 |  | 496,217,290 |  |
|  | Ratio of Ns (%) | 6.98 |  |  |  | 6.71 |  |  |  |
|  | GC content (%) | 36.52 |  |  |  | 36.58 |  |  |  |
| Chromosome Level |  | 461,198,141 | 31 |  |  | 435,876,255 | 31 |  |  |
| Chromosome Level (%) |  | 84.83 |  |  |  | 81.94 |  |  |  |

### 3.2 Evaluation of the assembly

The GC content for the genomes of the male and female individual was found to be 35.52% and 36.57% (Table 2), respectively, which showed a similar level with most closely related Lepidoptera species ranging from 31.6% to 37.7%^61^. We used the Benchmarking Universal Single-Copy Orthologs (BUSCO: version 2.0)^62^ and Core Eukaryotic Genes (CEGMA)^63^ to evaluated the completeness of the two assemblies. In BUSCO analysis, genomes of the male and female samples covered 95% and 94.5% complete BUSCO genes (Table S3). In the CEGMA analysis, 83.47% and 85.48% complete core eukaryotic genes were found for the two genomes (Table S4). This is better than all FAW genomes that has been published (PRJNA380964; PRJNA257248; PRJEB13110; and PRJNA344686). Besides, we also mapped the sequencing data generated from libraries of Hi-C, MatePair5K, WGS, and the RNA-seq to the assembled male genome. The mapping rates were all higher than 90% (Table 3), and the insert size were also consistent with the libraries, except for the MatePair5K, probably because the large insert size cannot ensure that the one pair reads they mapped to the same scaffold. EST sequences of the FAW were downloaded from NCBI and the transcripts were assembled without reference. We mapped these EST sequences and transcripts to the male reference genome we assembled, and the results showed that more than 90% EST sequences and more than 80% transcripts we assembled could be found on the assembled male genome (Table S5). However, it is noteworthy that the transcript from SFgdRNA 2 has a lower mapping rate than that of SFgdRNA 1. We inferred that this resulted due to the genetic differences between the C strain and the R strain, because the SFgdRNA 2 sample was identified belong to the R strain. Overall, all the above results well testified the completeness of the two genomes.

**Table 3.** Mapping reads against the assembled male genome using raw reads generated by different libraries

| Type | HIC | MatePair | stLFR | RNA-seq | WGS |
| --- | --- | --- | --- | --- | --- |
| Total Mapped Reads | 93.58% | 93.68% | 95.60% | 98.98% | 90.71% |
| Perfect Match | 38.81% | 38.54% | 43.67% | 52.14% | 27.76% |
| Unique Match | 76.39% | 77.15% | 83.23% | 76.99% | 74.77% |
| Total Unmapped Reads | 6.42% | 6.32% | 4.40% | 1.02% | 9.29% |
| Total FullMapped Reads | 32.48% | 30.06% | 71.45% | 52.46% | 49.40% |

### 3.3 Annotation

We firstly used Repeat Modeler (v1.0.11), LTR finder (v1.0.5) and repeatscount (v1.0.5) methods to identify *de novo* repeat motifs by modeling *ab initio*, and these repeat motifs were added into the RepBase^64^ library as known repeat elements. We then performed the RepeatMasker^65^ to mask the assembly, using the combined RepBase library. Usually, Repeat elements take a substantial part of the genome and contribute as important events to genome evolution^61,66,67^. In this study, by the combination of *de novo* and homology-based searching, 153 Mbp repeat elements were finally identified for the male FAW, and accounting for 28.24% the FAW genomes.

Gene prediction was carried out by both the homology-based and *de novo* methods using repeat masked genomes. For the *de novo* prediction, we used Augustus, glimmerHMM and SNAP (Table S6). For the homology-based approaches, *Bombyx mori*, *Danaus plexippus*, *Drosophila melanogaster* and *Spodoptera litura* genomes were used for homology alignments using the TblastN. Moreover, transcripts that were predicted with RNA-seq, Gene sets were then merged to form a non-redundant gene set with GLEAN; then all annotated genes were checked and filtered manually. A total of 22201 genes were finally obtained for the male samples (Table S6).

In the final gene set we identified, we found 94.2% compete for BUSCO genes and 95.16% CEGMA genes, which were all better than the published FAW genome (Table S7, Table S8). Of these identified genes, 93.48% was confirmed that have functions (Table S9), which was facilitated the further exploration of the functions. Besides, we also found 60 miRNAs, 840 tRNAs and 197 rRNAs by using the homology prediction method.

### 3.4 The transcriptome analysis of the larvae

After filtering, we finally obtained 58Gb clean data with 341,526,489 cleaned reads. These reads were assembled into 72,604 contigs with the N50 of 2077bp. These contigs were further assembled into scaffolds, and the scaffolds were further assembled to 51,495 unigenes by clustering and removing redundancy. The contig number in our study are significantly higher than that in the study of Kakumani *et al^68^*. This maybe result from that we only used a single method for assembly. We also calculated the expression abundance for unigenes between the fifth-instar and sixth-instar larvae. The result showed 2,648 differentially expressed genes (DEGs). We further performed the clustering analysis to cluster genes with identical or similar expressed behaviors. Remarkable expression difference was found between the fifth-instar larvae and the sixth-instar larvae (Figure S1). This difference was in consistent with the different strains of the two larvae (we described in 3.6). However, if the different instar contributes to the differential expression, it was further confirmed by more detailed analysis.

### 3.5 Comparison to other published lepidopteran genomes

To further explore the detailed relationship between the FAW and its other lepidopteran relatives. We constructed a phylogenetic tree of nine genomes using 2,001 single copy genes downloaded from NCBI and insectbase (Table S10). The result showed that the *S. frugiperda* which we sequenced actually clustered with its most related species *S. litura*, which is in accordance with the study by Cheng *et al* ^69^(Figure 3).

**Figure 3.**
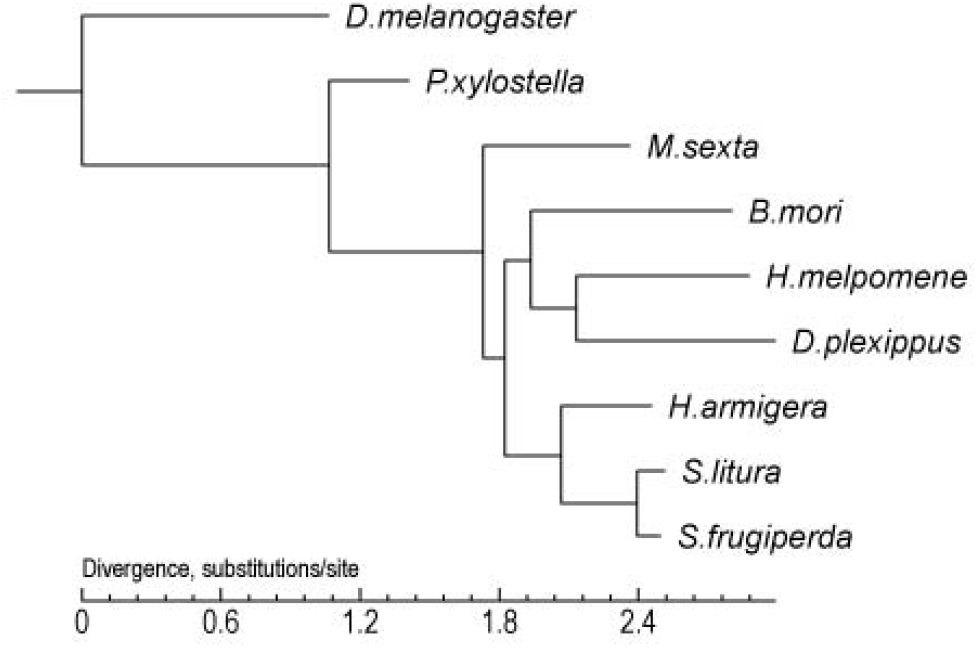
The phylogenetic relationships among nine lepidopteran genomes.

Through the gene family analysis, a total of 12,516 gene families were found in the *S. frugiperda* genome, including 20,012 genes. Of these gene families, 324 are specific to the *S. frugiperda* compared to the other seven species (Table S11). Then, we analyzed the 34 functional gene families of insects, finding some expanded gene families, including cytochrome p450, glutathione s-transferase, and hydrolase (Table 4). The cytochrome p450 gene family is closely related with intensified detoxification^69^, the genes in this family of the *S. frugiperda* is 200, more than that of *S. litura*, which indicated that the *S. frugiperda* was more polyphagous than *S. litura*. This is also consistent with the habits of *S. frugiperda*. The expanded glutathione s-transferase gene family was proved that could enhance the insecticides tolerance of the *S. litura^69^*. In this study, we found more genes for *S. frugiperda*, which indicated that the *S. frugiperda* was probably easier to gain resistance to pesticides. These gene families are a valuable genetic source to develop more effective pesticides or other methods to manage the FAW.

**Table 4.** Identified genes in gene families of 9 insects

|  | DME | PXY | MSE | BMO | HME | DPL | HAR | SLI | SFR |
| --- | --- | --- | --- | --- | --- | --- | --- | --- | --- |
| ABC transporter | 55 | 76 | 40 | 40 | 35 | 38 | 36 | 86 | 66 |
| acetylcholine receptor | 18 | 27 | 20 | 27 | 24 | 28 | 22 | 22 | 27 |
| Acetylcholinesterase | 27 | 39 | 62 | 51 | 32 | 27 | 82 | 87 | 46 |
| alkaline phosphatase | 13 | 10 | 11 | 10 | 8 | 8 | 8 | 11 | 12 |
| aminopeptidase | 45 | 53 | 49 | 37 | 35 | 39 | 15 | 19 | 24 |
| carboxylesterase | 30 | 55 | 90 | 70 | 50 | 54 | 100 | 110 | 84 |
| Chitinase | 45 | 27 | 28 | 24 | 26 | 32 | 23 | 26 | 46 |
| chloride channel | 16 | 25 | 20 | 23 | 18 | 19 | 19 | 16 | 24 |
| CTL | 2 | 4 | 3 | 2 | 2 | 2 | 3 | 4 | 4 |
| cytochrome p450 | 104 | 108 | 144 | 96 | 130 | 99 | 121 | 132 | 200 |
| DNA methyltransferase | 1 | 1 | 1 | 1 | 1 | 1 | 1 | 1 | 1 |
| ecdysone receptor | 1 | 1 | 3 | 3 | 3 | 2 | 3 | 1 | 3 |
| GABA | 15 | 16 | 32 | 9 | 12 | 15 | 16 | 14 | 10 |
| glutamate-gated chloride channel | 10 | 14 | 12 | 13 | 10 | 10 | 12 | 12 | 14 |
| Glutathione s-transferase | 50 | 28 | 42 | 25 | 23 | 25 | 46 | 47 | 60 |
| glycosyltransferase | 65 | 25 | 23 | 19 | 24 | 22 | 19 | 21 | 31 |
| G protein | 23 | 42 | 36 | 31 | 26 | 29 | 29 | 29 | 30 |
| gustatory receptor | 54 | 69 | 45 | 76 | 73 | 68 | 197 | 238 | 220 |
| heat shock protein | 45 | 38 | 54 | 35 | 29 | 46 | 40 | 48 | 45 |
| hydrolase | 213 | 250 | 243 | 209 | 181 | 204 | 267 | 310 | 396 |
| immunoglobulin | 65 | 51 | 70 | 76 | 56 | 69 | 71 | 67 | 80 |
| odorant-binding protein | 62 | 65 | 70 | 51 | 87 | 63 | 77 | 61 | 70 |
| odorant receptor | 67 | 82 | 79 | 43 | 73 | 64 | 87 | 75 | 75 |
| Painless | 1 | 1 | 1 | 1 | 1 | 1 | 1 | 1 | 1 |
| pheromone | 43 | 22 | 29 | 15 | 19 | 14 | 16 | 14 | 19 |
| protease inhibitor | 75 | 33 | 60 | 38 | 26 | 36 | 46 | 39 | 51 |
| Ryanodine receptor | 6 | 12 | 3 | 7 | 4 | 8 | 6 | 3 | 6 |
| sensory neuron membrane protein | 3 | 5 | 4 | 5 | 4 | 7 | 7 | 3 | 10 |
| serpin | 35 | 23 | 30 | 18 | 19 | 24 | 21 | 18 | 26 |
| sirtuin | 6 | 4 | 3 | 5 | 5 | 5 | 7 | 7 | 7 |
| sodium channel | 38 | 30 | 33 | 24 | 15 | 27 | 30 | 25 | 36 |
| sugar transporter | 17 | 51 | 42 | 36 | 40 | 37 | 41 | 40 | 59 |
| superoxide dismutase | 17 | 18 | 12 | 12 | 9 | 11 | 12 | 13 | 36 |
| Vitellogenin receptor | 8 | 14 | 15 | 11 | 9 | 12 | 16 | 14 | 15 |

Note: gene families come from http://www.insect-genome.com/genefamily/gene-family.php

### 3.6 FAW in China includes both the C and R strain, possibly invaded from Africa

There are three strain-specific sites (E4165, E4168 and E4183) in the fourth exon of the *Tpi* gene that can identify the C strain from the R strain for Western Hemisphere populations^57^. Especially, the E4183 is an effective diagnostic marker for *Tpi* gene to define the C or R strain. In this study, two Yunnan samples and one Guangdong sample were identified as the C strain and the other one Guangdong sample was identified as the R strain (Figure 4). The phylogenetic tree showed that the two Yunnan samples were clustered in the clade that consisted of all C strain individuals, which strengthened the results inferred using the strain-specific sites in the fourth exon (Figure 5). This result at least showed that the FAW invaded into China included both the strains. However, the detailed population genetic structure and the frequencies of the two strains in the Chinese population need more information from the population-level studies.

**Figure 4.**
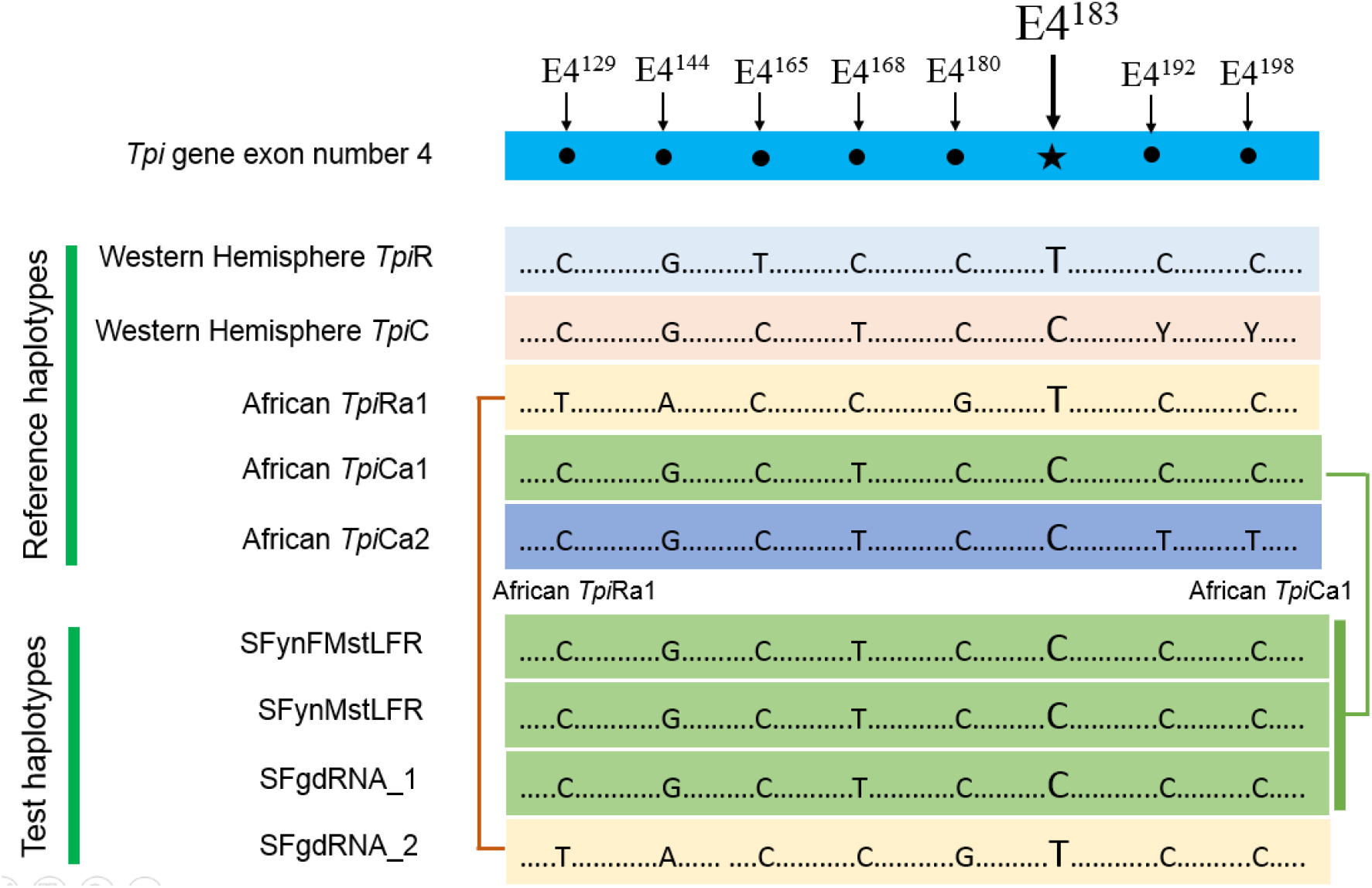
The identification of C and R strains.

**Figure 5.**
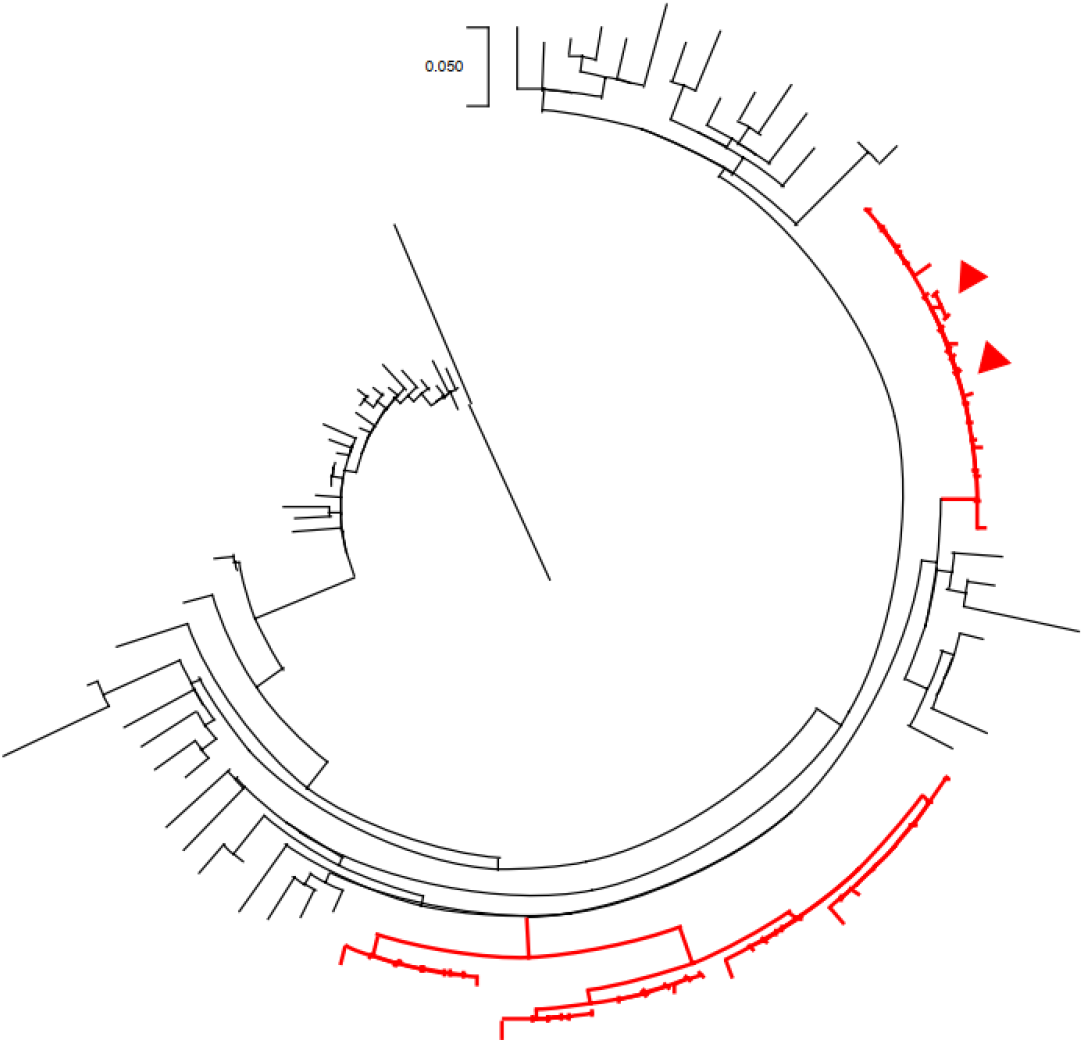
The phylogenetic tree to identify the FAW strains of collected from Yunnan, China. The red branched presented the C strain, the red triangles present the two Yunnan samples.

To further confirm the possible source of the Chinese FAW, we compared the haplotype consisted of all eight polymorphisms in the fourth exon of the *Tpi* gene as shown in the Figure 4. We found that all the four individuals hold identical haplotype with the African population, including three *Tpi*Ca1 and one *Tpi*Ra1. Although the haplotype of C strain was shared by the African and the Western Hemisphere populations^57^, the *Tpi*Ra1 has not been detected in any Western Hemisphere populations^57^ showing the uniqueness to African populations. The finding of the *Tpi*Ra1 haplotype in the Guangdong population indicated that there were at least parts of the FAW populations in China that was invaded from Africa, probably through the frequent commercial trade and passenger transportation between Africa and China^5^. However, we cannot confirm other sources because of the small sample size we used here. The strain information and possible invasion source found in this study will be extremely important for making effective strategies to manage the FAWs in China.

## 4 Conclusions

In summary, we assembled two chromosome scale genomes of the fall armyworm, representing one male and one female individual procured from Yunnan province of China. The genome sizes were identified as 542.42 Mb with N50 of 14.16 Mb, and 530.77 Mb with N50 of 14.89 Mb for the male and female FAW, respectively.. The completeness of the two genomes are better than all previously published FAW genomes which is evident by the BUSCO and CEGMA analysis. A total of 22,201 genes were predicted in the male genome, and 12,516 gene families were found in the *S. frugiperda* genome, including 20,012 genes. Of these gene families, we found expansion of cytochrome p450 and glutathione s-transferase gene families, which were closely related to the function of intensified detoxification and pesticides tolerance. We finally identified both the R strain and C strain individuals in the Chinese population, showed that the Chinese FAW was most likely invaded from Africa. The strain information and possible invasion source found in this study will be extremely important for making the effective strategies to manage the FAWs in China.

## Acknowledgements

This study was supported by the Guangdong Provincial Key Laboratory of core colletion of corp genetic resources research and application (NO.2011A091000047), Guangdong Provincial Key Laboratory of Genome Read and Write (grant no. 2017B030301011) and Key Laboratory of Genomics, Ministry of Agriculture.

## Data availability

The Raw sequencing data and the two chromosome level genome assemblies have been deposited to the CNSA (CNGB Nucleotide Sequence Archive) with accession CNP0000513 (https://db.cngb.org/cnsa/).

## Author Contributions

Huanming Yang, Le Kang, Jun Sheng, Youyong Zhu, Yang Dong, Xin Liu and Huan Liu designed the research. Dongming Fang, Tianming Lan and Yue Chang, Hongli Wang, Fangneng Huang, Wei Dong and Guangyi Fan performed the data analysis. Xiaofang Chen, Haorong Lu, Ping Liu, Tongxian Liu, Rushi Hao, Bin Chen, Shusheng Zhu, Zhihui Lu and Haimeng Li performed the DNA and RNA extraction and the library preparation. Huan Liu, Tianming Lan, Yang Dong, Wei Guo, Shuqi He, Le Chen and Lihua Lyu collected the samples. Tianming Lan, Dongming Fang, Hongli Wang, Sunil Kumar Sahu and Furong Gui wrote and revised the manuscript. All the authors read and revised the final version of the manuscript.

