## Supplementary figures and images for "Chromosome level draft genomes of the fall armyworm, *Spodoptera frugiperda* (Lepidoptera: Noctuidae), an alien invasive pest in China"

### Figure S1

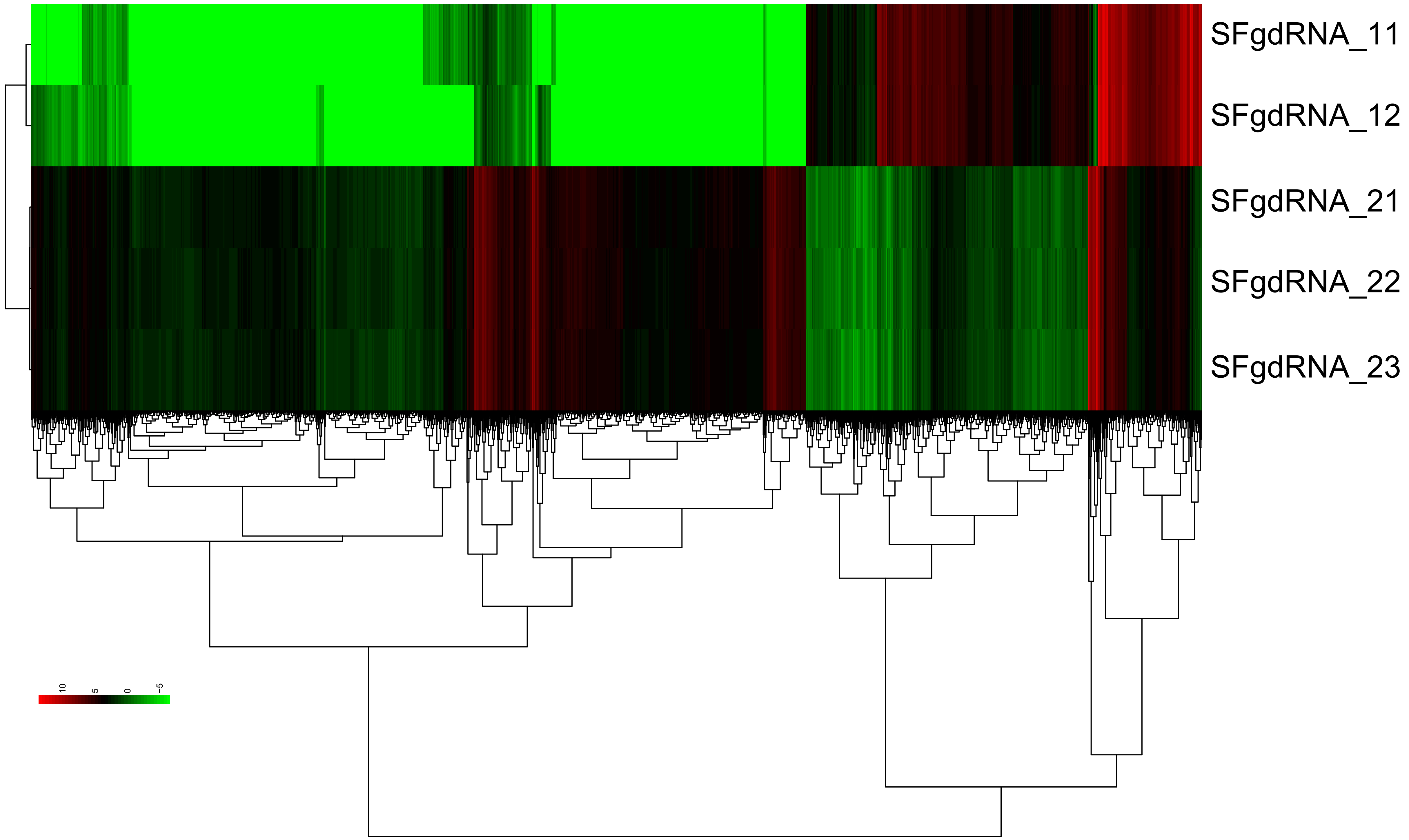
